# JMJD3 Deficiency in Midbrain Dopamine Neurons Disturbs Dopamine Synthesis and Aggravates Chronic Inflammatory Pain

**DOI:** 10.1101/2022.12.01.518657

**Authors:** Xi-Biao He, Fang Guo, Wei Zhang, Weidong Le, Yong Zheng, Yongjun Ma, Sang-Hun Lee, Hui-Jing Wang, Yi Wu, Qinming Zhou

**Author notes:** Corresponding author Xi-Biao He, Address: 279 Zhouzhu Highway, Pudong New Area, Shanghai, 201318, China. These authors contribute equally to this work (Fang Guo, Wei Zhang). Institute of Neuroscience, State Key Laboratory of Neuroscience, CAS Center for Excellence in Brain Science and Intelligence Technology, Chinese Academy of Sciences, Shanghai 200031, China.

## Abstract

Chronic pain is associated with midbrain dopamine levels. The molecules and mechanisms modulating this association remain to be elucidated. By using conditional knockout mice, we report that JMJD3 deficiency in midbrain dopamine neurons causes prolonged mechanical hyperalgesia regardless of sex and age. Both genetic defect and pharmaceutical inhibition of JMJD3 decrease dopamine level in midbrain and striatum, resulting from reduced tyrosine hydroxylase expression in midbrain dopamine neurons. Furthermore, epigenetic experiments reveal that JMJD3 is indispensable for the transcription of tyrosine hydroxylase through direct and indirect manners. These findings suggest JMJD3 and midbrain dopamine neurons as novel players and pharmaceutical targets for chronic pain regulation.

## Introduction

Dopamine (DA) neurotransmission is widely implicated in the modulation of pain (Ossipov et al., 2010; Taylor et al., 2016). Midbrain DA neurons of the ventral tegmental area (VTA) and substantia nigra pars compacta (SNc) are the major cell types that produce DA in the central nervous system. Increasing evidence indicate a strong association between chronic pain and decreased DA activities in midbrain DA neurons (Yang et al., 2021; Markovic et al., 2021; Huang et al., 2020; Martikainen et al., 2015; Scott et al., 2006; Hnasko et al., 2005), whereas the regulatory molecules and underlying mechanisms are much less understood.

Tyrosine hydroxylase (TH) is the rate-limiting enzyme for DA biosynthesis, which is down-regulated in pathological midbrain DA neurons in Parkinson’s disease (PD) (Johnson et al., 2018). Although chronic pain is a common non-motor symptom in PD patients (Beiske et al., 2009), whether decreased TH activity plays a causal role in PD-related pain hyperactivity remains unclear. Depending on cellular context, TH expression and activity are epigenetically, transcriptionally and post-translationally regulated (Lenartowski and Goc, 2011). Despite of a large number of transcription factors identified including the master regulator nuclear receptor related 1 (NURR1) (van Heesbeen et al., 2013), the epigenetic factors that affect TH-mediated DA synthesis are just beginning to get revealed.

Jumonji C-domain-containing histone deacetylase 3 (JMJD3) is a histone deacetylase specifically demethylating histone 3 lysine 27 (H3K27), which is related to chromatin remodeling that switches transcription from repression to activation (Hong et al., 2007, 3). JMJD3 participates in a wide range of biological processes in various types of cells, from early embryonic development to cell senescence (Burchfield et al., 2015). However, little is known about the roles of JMJD3 in midbrain DA neurons, except for our previous study reporting it as an indispensable transcription factor for the DA phenotype commitment of prenatal DA progenitor cells (He et al., 2015).

To understand the role of JMJD3 in adult midbrain DA neurons, we have constructed DA neuron-specific JMJD3 conditional knockout (cKO) mice and investigated how JMJD3 deficiency affects midbrain DA neurons on molecular, cellular and behavioral levels.

## Results & Discussion

### *Th*-specific *Jmjd3* knockout exaggerates chronic inflammatory pain

We created a heterogeneous cKO mouse model by crossbreeding transgenic mice carrying *Th-*CRE and loxP-floxed *Jmjd3* loci, respectively. Offspring mice carrying *Th-*CRE and one allele of loxP-floxed *Jmjd3* were used as cKO and those carrying *Th-*CRE and normal *Jmjd3* as control (WT; Supplemental Fig. S1A). To understand the behavioral phenotype of this model, WT and cKO mice in different sexes and ages underwent inflammation-induced mechanical hyperalgesia test and accelerating rotarod test respectively, to determine their pain sensitivity and locomotor coordination, both of which are midbrain DA-related.

To induce mechanical hyperalgesia, complete Freund’s adjuvant (CFA) was injected intraplantarly in both WT and cKO mice. Pain sensitivity was measured at various time points over 18 days by Von Frey’s hairs. As expected, WT and cKO mice showed no difference in basal nociception. CFA caused significant pain hypersensitivity over the first three hours, regardless of sex and age, indicating an intact perception of acute pain for this mouse model (Fig. 1A-D). The pain sensitivity was not attenuated until the eighth day (D8) in both male and female young WT mice, followed by a nearly complete recovery from D13, validating the confidence of this test. By contrast, the hyperalgesia lasted for at least 13 days in young cKO mice of both sexes, and was only slightly attenuated at D18 (Fig. 1A, B). This result indicated a sex-independent prolonged pain hypersensitivity in young cKO mice. On the other hand, the hyperalgesia was not attenuated in old male WT mice, and to a much milder extent in old female ones over 18 days (Fig. 1C, D), suggesting that the pain maintenance was worsened along aging. In comparison, similar to young cKO mice, neither old male nor female cKO mice showed attenuation of hyperalgesia over 18 days. These results demonstrated that *Jmjd3* cKO caused age- and sex-independent pain maintenance. To confirm the pain specificity, the thermal hyperalgesia was measured in parallel using a Hargreaves method (Hargreaves et al., 1988) and no difference was observed (Supplemental Fig. S1B).

**Figure 1.**
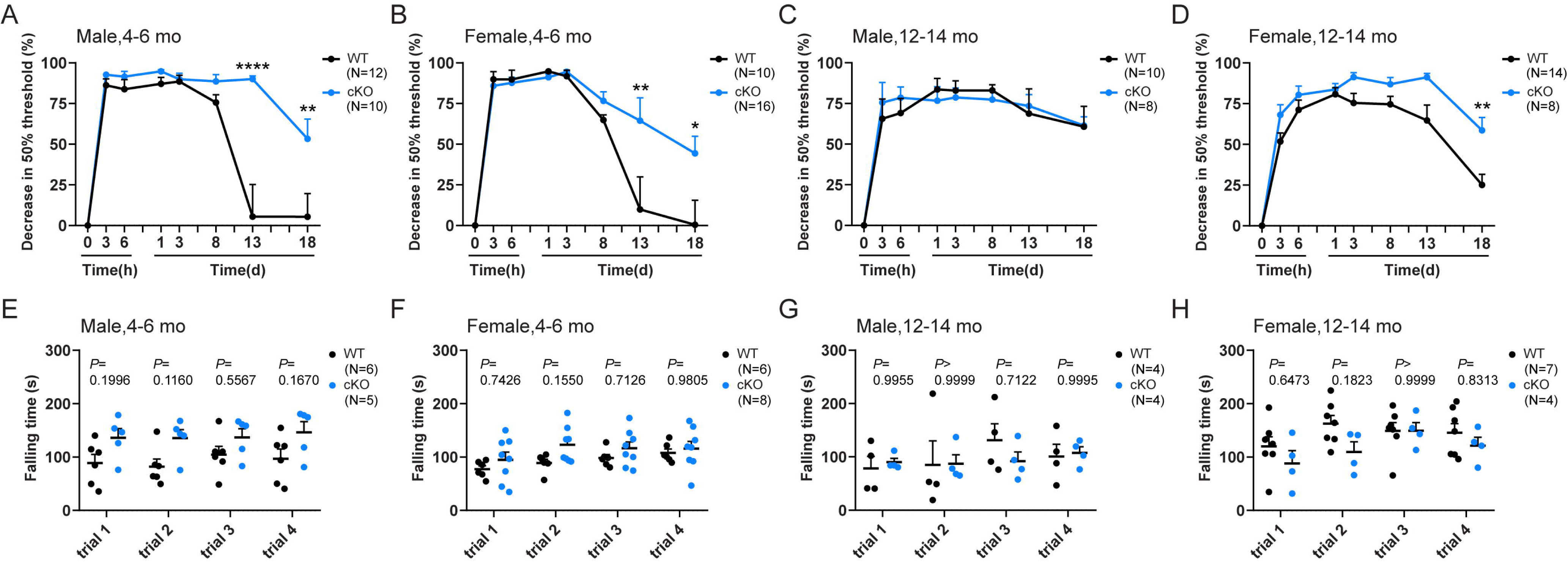
Tyrosine hydroxylase-specific JMJD3 knockout affects inflammation-induced mechanical hyperalgesia but not locomotor functions. (A-D) Conditional knockout (cKO) mice and their control littermates (WT) received an intraplantar injection of complete Freund’s adjuvant and changes in 50% paw withdrawal threshold assessed by Von Frey’s hair were monitored over 18 days. Mice were grouped into young (4-6 months old (mo); A, B) and old (12-14 mo; C, D), male (A, C) and female (B, D). (E-H) In parallel, mice underwent the accelerating rotarod test daily for four consecutive days. The falling time of each mouse was averaged of two repeats. Data represent mean ± SEM. \**P* < 0.05, \*\**P* < 0.01, \*\*\*\**P* < 0.0001; two-way ANOVA with Sidak’s multiple comparisons test.

The accelerating rotarod test is often used to assess locomotor coordination in rodents. In addition, the test can be modified for four consecutive days to enable a further demonstration of motor skill learning, which more likely reflects the vulnerability of DA depletion (Chagniel et al., 2012; Shiotsuki et al., 2010). As previously reported (Shoji et al., 2016), a decrease of locomotor activity was noticed in old male mice in comparison to young male ones, validating the specificity of the test. However, no significant difference of falling latency was found between WT and cKO mice within four trials, regardless of sex and age (Fig. 1E-H). In addition, no difference of grip strength was found (Supplemental Fig. S1C). Taken together, these results suggest that *Jmjd3* cKO specifically affects chronic pain maintenance rather than locomotor coordination.

### DA contents are decreased in cKO mouse midbrain and striatum

To understand the biological alterations in JMJD3-deficient mouse brain, the levels of two catecholamines DA and norepinephrine (NE) were measured by high performance liquid chromatography-mass spectrometry (HPLC-MS) in several TH- enriched brain regions of young WT and cKO mice. The results demonstrated that the levels of DA in the midbrain and striatum of cKO mice were significantly lower than those of WT mice in both sexes, whereas no significant difference was observed in frontal cortex, cerebellum and brainstem (Fig. 2A and Supplemental Fig. S2A). By contrast, the level of NE was not altered in any brain regions examined (Fig. 2B and Supplemental Fig. S2B), suggesting that the *Jmjd3* cKO effect was confined to midbrain DA system.

**Figure 2.**
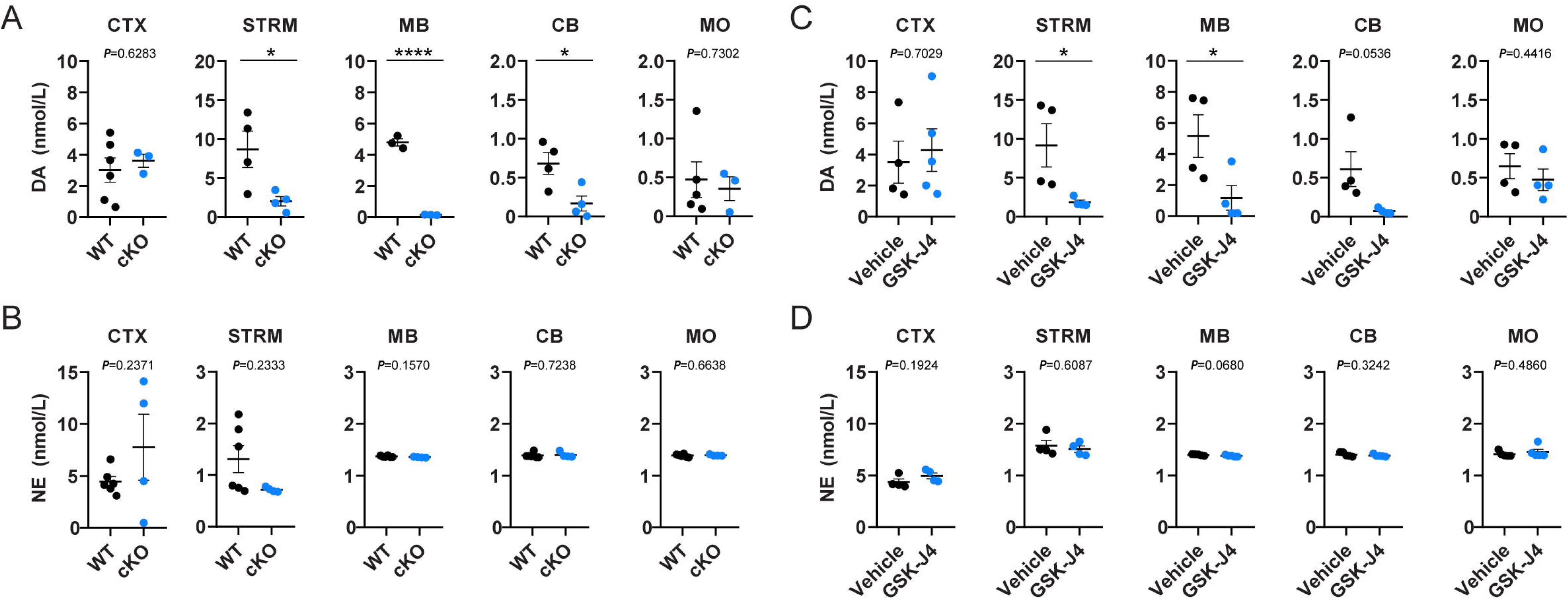
Dopamine levels are decreased in the midbrain and striatum of two mouse models of JMJD3 deficiency. Brain tissues from frontal cortex (CTX), striatum (STRM), midbrain (MB), cerebellum (CB) and medulla oblongata (MO) were micro-dissected, homogenized and the contents of dopamine (DA) and norepinephrine (NE) were measured by high performance liquid chromatography-mass spectrometry. (A, B) 4-month-old male conditional knockout (cKO) mice and their control littermates (WT). (C, D) 4-month-old male C57BL/6 mice underwent 5 consecutive days of intra-peritoneal injection of GSK-J4 (20mg/kg daily). In parallel, dimethyl sulfoxide was injected as Vehicle. N= 4-7 mice. Data represent mean ± SEM. \**P* < 0.05, \*\*\*\**P* < 0.0001; Student’s t-test.

To confirm the results above, a brain-penetrating JMJD3 inhibitor GSK-J4 (Kruidenier et al., 2012) was daily injected intra-peritoneally into young male mice for five days, following by the same experimental procedure of HPLC-MS. In line with results from genetic defect mouse model, decreased levels of DA but not NE were specifically observed in the midbrain and striatum of GSK-J4-injected mouse brains (Fig. 2C, D). These results provide consistent evidence supporting the vulnerability of midbrain DA contents to JMJD3 activity.

### TH and NURR1 expressions are decreased in midbrain DA neurons of cKO and GSK-J4 mice

The biochemical findings above suggested that the DA biosynthesis in midbrain DA neurons might be affected by JMJD3. Thus, we examined the TH protein expression in the midbrains of young male WT and cKO mice by immunofluorescence microscopy analysis. We found that both the TH immunoreactivity and cell number of TH-positive DA neurons were significantly lower in the SNc and VTA of cKO mice in comparison to WT (Fig. 3A, B). Similarly, TH immunoreactivity was also decreased in the SNc and VTA of GSK-J4 mice, whereas no significant difference of total TH-positive cell number in either region was observed (Fig. 3C, D). It is worth mentioning that the decrease of TH-positive cell number is more likely resulted from the attenuated TH expression rather than the loss of DA neurons *per se*, as neither the total amount of cells nor the overall morphology in midbrain seems not altered (data not shown). These results suggested that the down-regulation of TH expression in midbrain DA neurons might underlie JMJD3 deficiency-induced loss of DA contents.

**Figure 3.**
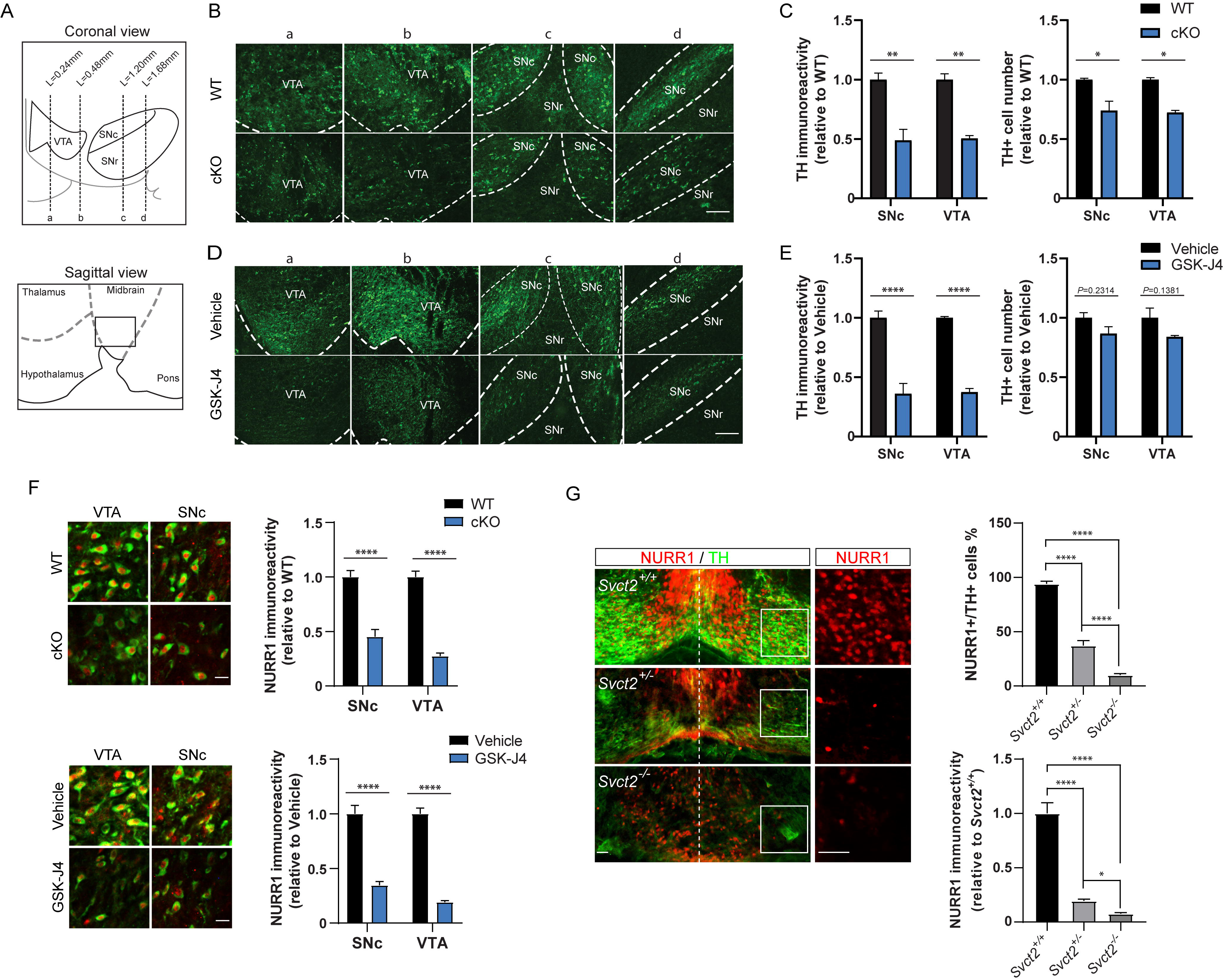
Decreased tyrosine hydroxylase and NURR1 expression in midbrain dopamine neurons of two JMJD3 deficiency models. (A) Schematics of mouse midbrain compromised of ventral tegmental area (VTA), substantia nigra pars compacta (SNc) and pars reticulata (SNr). Dotted lines a-d in coronal view and box in sagittal view indicate positions shown in B and D. (B) Representative immunofluorescence images of tyrosine hydroxylase (TH)-positive midbrain dopamine (DA) neurons in heterogeneous conditional knockout (cKO) mice and their control littermates (WT). Scale bar represents 100 μm. (C) Quantification of TH immunoreactivity and cell number of DA neurons in VTA and SNc in WT and cKO mice. (D) Representative immunofluorescence images of TH-positive DA neurons in GSK-J4 injected mice and their control (Vehicle). Scale bar represents 100 μm. (E) Quantification of TH immunoreactivity and cell number of DA neurons in VTA and SNc in GSK-J4 injected mice and control. (F) Representative immunofluorescence images and quantification of NURR1 immunoreactivity showing decreased NURR1 (red) expression in TH-positive DA neurons (green) in VTA and SNc of cKO and GSK-J4 mice. Scale bar represents 20 μm. (G) Representative immunofluorescence images and quantification showing effect of *Svct2* KO on TH and NURR1 expression in mouse embryonic day 14.5 ventral mesencephalon. Insets show NURR1 expression in higher magnification. Scale bar represents 50 μm. For immunoreactivity analysis, at least 50 TH-positive cells were measured and the mean was calculated for one mouse. N = 3-5 mice. Data represent mean ± SEM. \**P* < 0.05, \*\**P* < 0.01, \*\*\*\**P* < 0.0001; Student’s t-test.

Given the essential role of transcription factor NURR1 in the initiation and maintenance of TH expression in midbrain DA neurons (Zetterström, 1997; Saucedo-Cardenas et al., 1998; Kadkhodaei et al., 2009, 2013), we further investigated whether NURR1 expression was altered in both cKO and GSK-J4 mice. Surprisingly, either cKO or GSK-J4 caused remarkable decrease of NURR1 expression in VTA and SNc DA neurons (Fig. 3E, F). To confirm this result, we recruited a previously reported (He et al., 2015) mouse model of indirect JMJD3 inhibition in which JMJD3 activity is prohibited through genetic loss of sodium vitamin C transporter 2 (*Svct2*). Although *Svct2* KO resulted in agenesis of the majority of DA neurons, a few DA progenitors were still able to migrate and differentiate into DA neurons with reduced expression of TH in the mantle zone of embryonic ventral mesencephalon. In these cells, the proportion that expressed NURR1 and the NURR1 immunoreactivity were significantly lower in comparison to those of wildtype (Fig. 3G). These histological experiments together indicated that the disturbance of NURR1-mediated TH expression might underlie JMJD3 deficiency-induced loss of DA contents.

### JMJD3 directly and indirectly activates *Th* transcription in midbrain DA neurons

JMJD3 is an epigenetic transcription activator. Thus, the transcription of *Jmjd3*, *Th* and *Nurr1* genes in WT and cKO mice was examined by real-time PCR analysis. Compared to WT, all three genes were specifically down-regulated in the midbrain and striatum of JMJD3 cKO mice (Fig. 4A), suggesting that TH and NURR1 were transcriptionally regulated by JMJD3. To confirm this result in vitro, primary culture of mouse midbrain DA neurons was challenged with 1 μM GSK-J4 for 1 day. GSK-J4 caused remarkable decreases of TH-positive cell number and NURR1 expression (Fig. 4B), along with down-regulation of both *Th* and *Nurr1* genes (Fig. 4C). Similar results were seen in a GSK-J4-treated DA neuronal cell line which consistently expressing TH (Crawford et al., 1992) (Supplemental Fig. S3A, B). In addition, short hairpin RNA-mediated JMJD3 knockdown confirmed the specificity of GSK-J4 (Supplemental Fig. S3C, D). These experiments demonstrated a transcriptional regulation of NURR1-TH pathway by JMJD3 in midbrain DA neurons.

**Figure 4.**
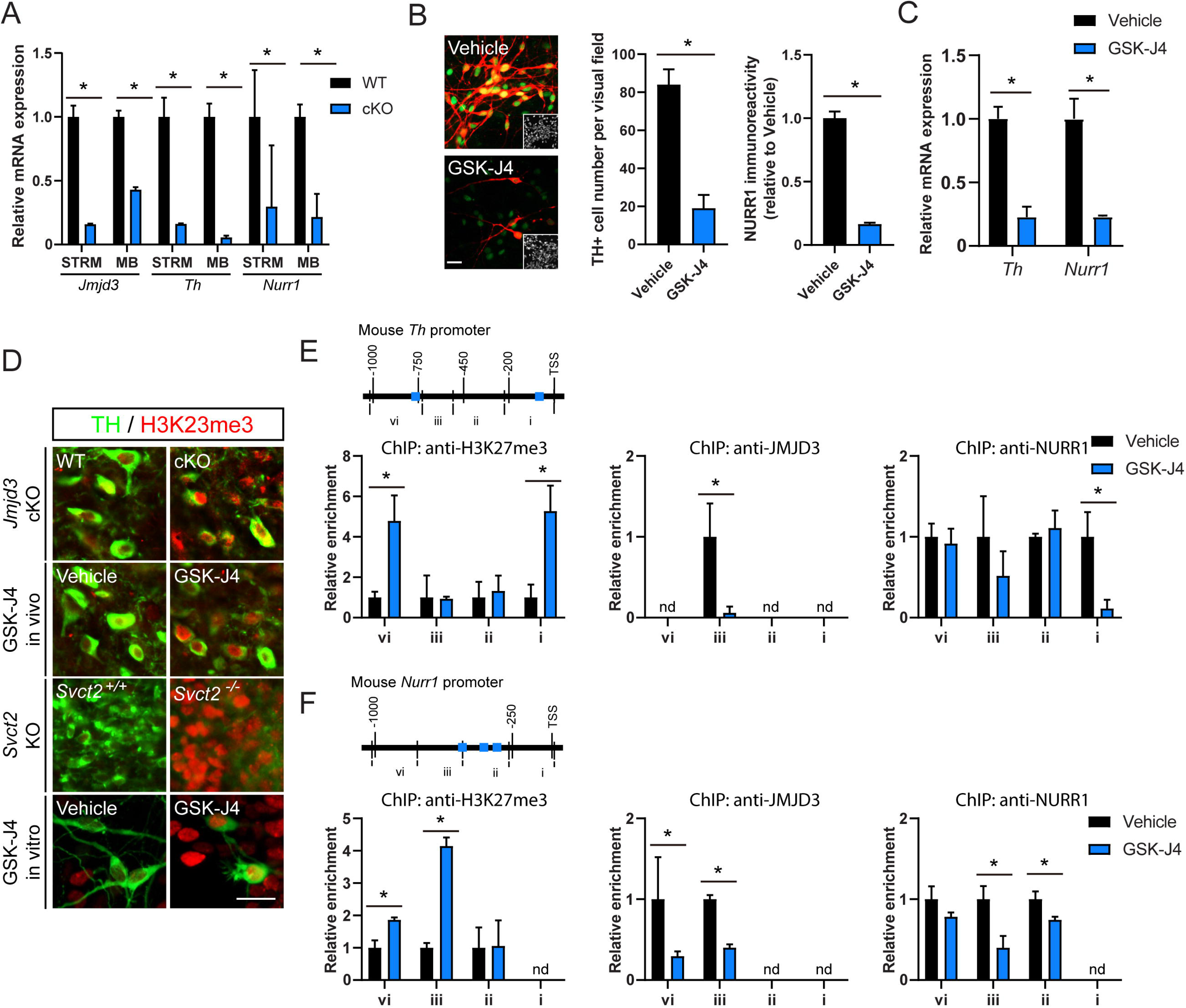
Epigenetic control of *Th* and *Nurr1* gene transcription by JMJD3. (A) Real-time PCR analysis of mRNA levels of *Jmjd3*, *Th* and *Nurr1* from striatum (STRM) and midbrain (MB) in conditional knockout mice (cKO) and their control littermates (WT). N = 3 mice. Data represent mean ± SEM. (B) Representative immunofluorescence images and quantification of NURR1 expression (green) in tyrosine hydroxylase (TH; red)-expressing dopamine (DA) neurons from primary mouse DA neuronal cell culture with or without treatment of GSK-J4. Insets show cell nuclei. Scale bar represents 20 μm. (C) Real-time PCR analysis of *Th* and *Nurr1* mRNA levels in primary DA neuronal culture after GSK-J4 treatment. (D) Representative immunofluorescence images of histone 3 lysine 27 tri-methylation (H3K27me3) in TH-positive midbrain DA neurons from cKO, GSK-J4 mice, *Svct2* KO mouse embryos and primary DA neuronal culture treated with GSK-J4. Scale bar represents 20 μm. (E, F) Chromatin immunoprecipitation (ChIP)-qPCR analysis of H3K27me3, JMJD3 and NURR1 enrichments on 1 kilo bp mouse *Th* (E) and *Nurr1* (F) promoters from primary DA neuronal cell samples with or without GSK-J4 treatment. Blue boxes represent consensus NURR1 binding sites. nd = not detected. N = 3 independent cell cultures. Data represent mean ± SEM. \**P* < 0.05; Student’s t-test and one-way ANOVA with Tukey’s post hoc test.

As a chromatin modifier, JMJD3 binds to DNA and specifically de-methylates H3K27 tri-methylation (H3K27me3). Indeed, global induction of H3K27me3 was observed in midbrain DA neurons from *Jmjd3* cKO, GSK-J4 mice and *Svct2* KO embryos, and in cultured DA neurons treated with GSK-J4 (Fig. 4D). By performing chromatin immunoprecipitation (ChIP) assays on primary culture and cell line of DA neurons, we found that H3K27me3 was specifically induced by GSK-J4 within 1 kilo bp of both *Th* and *Nurr1* gene promoters (Fig. 4E, F), suggesting a chromatin remodeling of both genes by JMJD3. This was further supported by region-specific decrease of JMJD3 binding induced by GSK-J4, within −745 to −475 bp of *Th* promoter, and −1091 to −510 bp of *Nurr1* promoter, respectively. Moreover, the enrichment of NURR1 was also greatly decreased within 0 to −203 bp of *Th* promoter and within −298 to −847 bp of *Nurr1* promoter (Fig. 4E, F), which correlated well with the locations of consensus NURR1 binding site as previously reported (Yi et al., 2014). Collectively, these results unraveled both *Nurr1* and *Th* promoters as direct target of JMJD3, and suggested a critical role of JMJD3-catylazed H3K27 de-methylation in facilitating NURR1-mediated transactivation of *Th* and likely *Nurr1* itself in midbrain DA neurons.

Our study report a previously unknown role of JMJD3 for the DA synthesis in adult mouse midbrain DA neurons, and establish a connection of this cell type-specific deficiency with chronic pain (illustrated in Fig. 5). Recent studies have reported chronic pain as an inducer of DA content decrease in midbrain particularly VTA DA neurons (Yang et al., 2021; Huang et al., 2020; Vergara et al., 2020). Evidence also exist suggesting that TH-expressing cells from other regions in central nervous system, such as A11 hypothalamic DA neurons and dorsal root ganglion neurons expressing C-low threshold mechanoreceptors (Brumovsky, 2016; Hnasko et al., 2005; Seal et al., 2009). However, we found no significant difference of the DA level and TH expression in hypothalamus between WT and cKO mice (data not shown), and JMJD3 expression is almost undetectable in TH-expressing dorsal root ganglion neurons (unpublished data). Therefore, we conclude that the contributions of these regions to JMJD3-related DA deficiency and hyperalgesia, if any at all, are not comparable to that of midbrain DA neurons.

**Figure 5.**
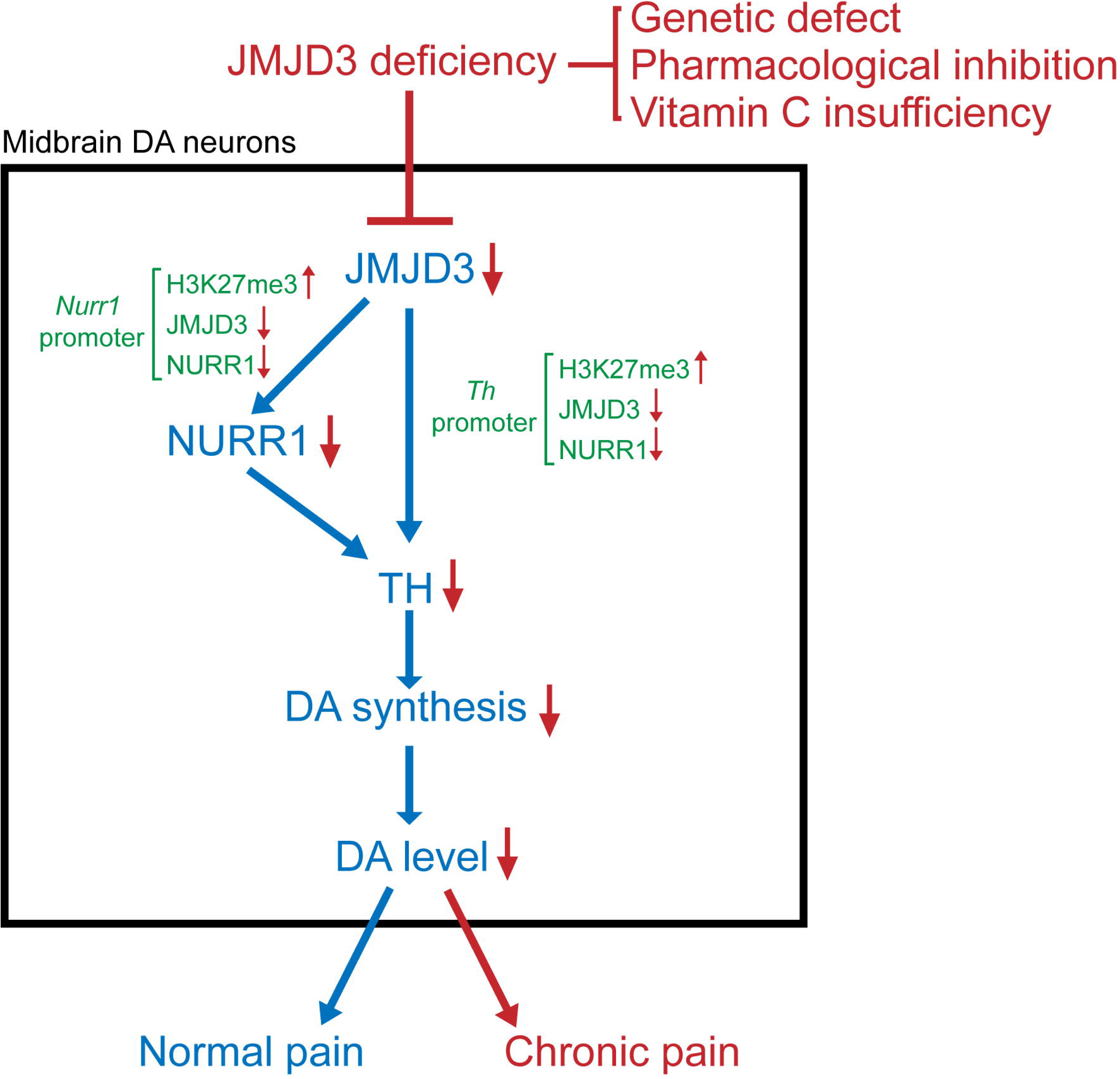
Schematics illustrating the association of JMJD3 deficiency in midbrain dopamine neurons and chronic pain. On behavioral level, conditional knockout (cKO) of *Jmjd3* in midbrain dopamine (DA) neurons in adult mouse causes prolonged inflammation-induced mechanical hyperalgesia. On tissue level, DA levels in striatum and midbrain are specifically decreased by cKO. On cell level, mRNA and protein expressions of DA synthesis rate-limiting enzyme tyrosine hydroxylase (TH) and its transcriptional master regulator nuclear receptor related 1 (NURR1) in midbrain DA neurons are decreased by cKO. On molecular level, cKO causes increase of repressive histone modification H3K27me3 and decreases of JMJD3 and NURR1 bindings on *Th* and *Nurr1* promoters, resulting in transcriptional inactivation of both genes. Therefore, JMJD3 deficiency in midbrain DA neurons induced by genetic, epigenetic and environmental causes is a novel risk factor for chronic pain.

Our finding reveals that the transcriptions of both TH and its master regulator NURR1 are under regulation of JMJD3 in a similar epigenetic manner, indicating that the DA content reduction is cell autonomous and multi-targeted. Importantly, midbrain DA neurons are not lost in cKO mice, demonstrating a non-toxic repression of TH and DA, which is distinguished from the neurotoxic characteristics commonly found in PD (Meredith et al., 2008). Another discrepancy is that the decreased TH activity in SNc does not cause motor abnormalities similar to those in PD, as evidenced by no difference between cKO and WT mice in the performance of accelerating rotarod. The physiological roles of JMJD3-mediated regulation of DA synthesis in normal midbrain DA neurons remain to be further studied.

A high consistency of neurotransmitter content changes is observed between drug-induced global JMJD3 inhibition and DA neuron-specific *Jmjd3* KO. Recent studies suggest promising potential of JMJD3 inhibitors in cancer therapy. For instance, GSK-J4 exhibits high efficiency in targeting pediatric brainstem glioma and high-risk neuroblastoma (Hashizume et al., 2014; Lochmann et al., 2018). However, our result shows that intra-peritoneal injection of GSK-J4 for 5 days would have remarkable impact on the metabolism of DA system in adult mouse brain. This might raise concerns about the adverse effects on DA-related functions and behaviors in pre-clinical and clinical trials with JMJD3 as therapeutic target.

## Materials and Methods

### Ethics

All animal experiments were approved by the animal ethics committee of Shanghai University of Medicine and Health Sciences and have been performed in accordance with the ethical standards laid down in the 1964 Declaration of Helsinki and its later amendments.

### Animals

For pharmaceutical JMJD3 inhibition experiments, 4-month-old male C57BL/6 mice were purchased from Shanghai Jiesijie Experimental Animal Co. GSK-J4 dissolved in dimethyl sulfoxide (DMSO) was daily injected intraperitoneally at 20 mg/kg body weight into mice for five consecutive days. In parallel, DMSO was injected in the same manner as control. For TH-specific JMJD3 cKO experiments, *Th*-CRE mice were crossbred with heterozygous floxed JMJD3 mice (a kind gift from Dr. Degui Charlie Chen) to generate cKO mice, whereas littermates of *Th*-CRE were used as WT. *Svct2* KO mouse embryos at embryonic day 14.5 were kindly provided by Dr. Fiona E. Harrison and have been described in our previous study (He et al., 2015). All animals were housed under standard conditions with controlled 12-hour light/dark cycles with food and water provided ad libitum.

### Cell culture

Primary midbrain DA neurons were cultured as previously described (He et al., 2015) with minor modifications. DA progenitors were extracted from embryonic day 11 ICR mouse ventral mesencephalon. At this stage, the SNc and VTA are not anatomically or molecularly distinct (Hu et al., 2004; La Manno et al., 2016), enabling these cellsdifferentiating into both DA subtypes. Briefly, progenitors were mechanically dissociated into single cells and plated onto 24-well plates or 6-well dishes pre-coated with 15 μg/ml poly-L-ornithine and 1 μg/ml fibronectin. Cells were cultured in serum-free Neurobasal media supplemented with N2 and B27 supplements and 1% Penecillin/Streptomycin for 7 days to reach a complete DA neuronal differentiation. After that, GSK-J4 or DMSO were treated for 24 hours. MES23.5 cells were cultured in DMEM supplemented with 10% fetal bovine serum and 1% Penecillin/Streptomycin. All cells were incubated in 5% CO_2_, 37 °C incubator.

### Real-time PCR analysis

RNAs were extracted from cultured cells and tissues using RNA extraction kit and reverse transcribed into cDNAs (both from TAKARA) according manufacturers’ instructions. Real-time PCR was performed on a Real-time system (Roche) using SYBR green supermix (Vazyme). The comparative cycle threshold method was used for quantification. Gene expression values were normalized to those of *Gapdh*. Primer sequences are listed in Supplemental Table S1A.

### Fluorescence immunostaining analysis

Brain tissues were pre-fixed with 4% paraformaldehyde by cardiac perfusion and post-fixed for 3 hours, dehydrated in 30% sucrose and stored in embedding compound (Tissue-Tek) at −70°C until cryosection. Tissues were cryosectioned into 14 μm thickness (Leica CM1950). Cultured cells were fixed with 4% paraformaldehyde for 20 mins. For immunostaining, tissue slices or cultured cells were permeabilized and blocked with PBS with 0.3% TritonX-100 and 1% bovine serum albumin for 40 mins, incubated with mouse anti-TH primary antibody (Sigma) diluted with blocking solution at 4°C overnight. Alexa Fluor series of second antibodies (Thermo Scientific) were applied for one hour at room temperature. Cells were finally counterstained with 4’,6-diamidino-2-phenylindole (DAPI) and examined using fluorescence microscope (Leica DMi8).

### HPLC-MS analysis

HPLC-MS experiments were performed as previously described (Pu et al., 2020; Sun et al., 2022). Briefly, samples from different brain regions after mouse sacrifice were homogenized in ice-cold 0.2M perchloric acid/methanol, sonicated (Bioruptor) and centrifuged at 10000 g for 20 mins. Supernatants were then filtered through 0.22 μm filter and stored in −80°C. At least four animals per genotype or drug treatment were used. Analytes were separated using a Model 1260 HPLC system consisting of a binary pump, on-line degasser, thermostated dual 54-well plate autosampler, and a thermosated column compartment (Agilent Technologies, Santa Clara, CA) with a Poroshell 120 SB-C18 reverse phase column (3.0x100 mm, 2.7 μm, particle size, Agilent Technologies, Santa Clara, CA) temperature-controlled at 25 LJ. The mobile phase consisted of A (1mM ammonium formate, 0.1%(v/v) formic acid and 1%(v/v) methanol in water) and B (1 mM ammonium formate 0.1%(v/v) formic acid and 0.9%(v/v) H2O in methanol). The gradient conditions of the mobile phase were as follows: 0 min, 5% B; 20 min,95% B; 21 min, 5% B and 28 min, 5% B. The flow rate of the mobile phase was 0.3 ml/min, and the injection volume was 10μl. For MS analysis, LC flow was coupled to an Agilent model 6470 triple quadrupole (QqQ) MS (Agilent Technologies, Santa Clara, CA), equipped with an Agilent Jet Stream (AJS) electrospray ionization (ESI) source. The mass spectrometer was operated with the following parameters: capillary voltage, 4000 V for positive and 3500 V for negative; nozzle voltage, 500 V. Nitrogen was used as the drying (5 L/min, 300 ℃), sheath (11 L/min, 250 ℃) and nebulizer gas (45 psi). The data was acquired using Agilent MassHunter Workstation Data Acquisition Software (revision B.04). Multiple-reaction-monitoring transitions parameters, such as precursor ions, product ions, fragmentor voltage, collision energy (CE) as well as polarity were shown in Supplemental Table S1. For the compounds which had two product ions, it was shown that the product ion was used as quantitative ion or qualitative one in Supplemental Table S1B. The dwell time was 50 ms for transitions.

### Mechanical hyperalgesia test

CFA-induced chronic mechanical hyperalgesia was assessed as previously described (Wang et al., 2013, 2). Briefly, baseline responses were measured using von Frey’s hairs (bending force range from 0.02-1.4 g, starting with 0.16g; Stoelting) after mice were placed on a wire grid without receiving any nociceptive stimulation for 2 hours. Next day, 20 μl CFA (Sigma-Aldrich) was injected intraplantarly, and the hyperalgesia responses were measured for various time points within 18 days. The hair force was increased or decreased according to the response, and the 50% paw withdrawal threshold was calculated using the up and-down method (Chaplan et al., 1994). Nociceptive responses include clear paw withdrawal, shaking, and licking. Stimulus and measurement were repeated if ambiguous responses such as ambulation occurred.

### Hargreaves test

Thermal algesia to noxious stimulus was measured using a hot/cold plate analgesia meter for mice (IITC) according to manufacturer’s instructions. mice were put on a glass plate and the hind paw plantar surface was exposed to a beam of radiant heat (intensity latency 8-12 s) according to the Hargreaves method (Hargreaves et al., 1988). Latencies of hind paw withdrawal were recorded. The heat stimulation was repeated 2 times at an interval of 5 min for each mouse and the mean of both paws was calculated.

### Grip strength test

Hindlimb grip strength was measured with a grip strength meter for mice consisted of a T-shaped metal bar connected to a force transducer (IITC). Briefly, mice were held by tails allowing them to grasp the metal bar with hind paws, followed by a pulling of the mice backwards by the tail until grip was lost. The peak force was automatically recorded in grams (g) by the device. The measurement of grip strength was repeated 5 times at an interval of 5 min for each mouse and the mean was calculated.

### Accelerating rotarod test

The test was performed on a rotarod apparatus (IITC) according to manufacturer’s manual. The 3 cm-diameter suspended rod was set to accelerate at a constant rate from 5 to 45 rpm in 240 s. Mice were pre-trained for 2 consecutive days before first trial. At each trial, mice were placed on the rod for a session of two repeats, with 180 s resting time allowed in between. A trial ended when the mouse fell off the rotarod or after reaching 240 s. The falling time was recorded and averaged of two repeats. Total four trials were performed for four consecutive days.

### Cell counting and statistics

For in vivo immunoreactivity analysis, at least 50 TH-positive cells were counted in each brain region for one animal. Data were derived from three mouse brains of same experimental group and expressed as mean ± SEM. For in vivo TH-positive cell number analysis, the total number of TH-positive cells from sagittal sections containing VTA and SNc in one mouse brain was counted. To compensate the double counting in adjacent regions, the Abercrombie correction factor was introduced as following: [N=n X (T/T+D)], in which N is the actual number of cells, n is the number of nuclear profiles, T is the section thickness, and D is the average diameter of nuclei. For in vitro analysis, cultured cells were counted in at least 10 random regions of each culture coverslip using an eyepiece grid at a magnification of 200 or 400X. Data are expressed as mean ± SEM of three independent cultures. For HPLC-MS analysis, data were derived from four to seven mouse brains of same transgenic background and expressed as mean ± SEM. Statistical comparisons were made using Student’s t-test, one-way ANOVA with Tukey’s post hoc analysis and two-way ANOVA with Sidak’s multiple comparisons test (Graphpad Prism).

## Acknowledgements

We thank Dr. Degui Charlie Chen and Dr. Fiona E. Harrison for providing *Jmjd3* loxP mice and *Svct2* KO embryos. X.B.H. is supported by the National Natural Science Foundation of China (31701287). F.G. is supported by the National Natural Science Foundation of China (32100779).

## Author Contributions

X.B.H. conceived the study, performed the experiments, generated figures and wrote the paper. F.G. performed the experiments, generated figures and drafted the manuscript. W.Z. performed HPLC-MS and drafted the manuscript. W.L. provided *Th*-CRE mice and provided administrative support. Y.Z., Y.M., Y.W. performed the experiments and generated figures. S.H.L. provided technical support for *Svct2* KO- related experiments. H.J.W. provided technical support for pain-related experiments. Q.Z. helped generate the mouse models. All authors have approved the final version of the manuscript.

## Supplemental Figure Legends

**Supplemental Figure S1.** (A) Genotyping of JMJD3 conditional knockout (cKO) mice. Genetic DNAs were extracted from tails of mice and applied to semi-quantitative PCR analysis to verify the loci carrying *Th*-CRE and *Jmjd3* loxP sequences using specific primers. Mice carrying one band by *Th*-CRE primer and two bands by loxP primer represent cKO and mice carrying one band by CRE primer and one band near 300 bp by loxP primer represent control genotype (WT). (B) Thermal hyperalgesia of cKO and WT mice in different sexes and ages were assessed by Hargreaves method. Latencies of hind paw withdrawal were recorded. (C) grip strength of cKO and WT mice in different sexes and ages were compared. Data represent mean ± SEM. Student’s t-test.

**Supplemental Figure S2.** Dopamine (DA) and norepinephrine (NE) levels were measured by high performance liquid chromatography-mass spectrometry in five brain regions including frontal cortex (CTX), striatum (STRM), midbrain (MB), cerebellum (CB) and medulla oblongata (MO) from 4-month-old female conditional knockout mice (cKO) and their control littermates (WT). N = 4-7 mice. Data represent mean ± SEM. \**P* < 0.05, Student’s t-test.

**Supplemental Figure S3.** (A) Representative immunofluorescence images and quantification of tyrosine hydroxylase (TH)-positive cells in MES23.5 DA neuronal cell culture with or without treatment of JMJD3 inhibitor GSK-J4. (B) Real-time PCR analysis of *Th* mRNA expression in response to different doses of GSK-J4 treatment. (C) Quantification of TH+ cell number in primary mouse DA neuronal cell culture underwent short hairpin RNA-mediated JMJD3 knockdown (sh-JMJD3) or control (plko.1). (D) Real-time PCR analysis confirmed that mRNA levels of *Jmjd3*, *Utx* and *Th* were affected by sh-JMJD3. Cell numbers were counted in 10 random areas of each culture coverslip using an eyepiece grid at a magnification ×100. Scale bar represents 100 μm. N = 3 independent cell cultures. Data represent mean ± SEM. \**P* < 0.05; one-way ANOVA with Tukey’s post hoc test.

