## Supplementary figures and images for "JMJD3 Deficiency in Midbrain Dopamine Neurons Disturbs Dopamine Synthesis and Aggravates Chronic Inflammatory Pain"

### Supplemental Figure 1

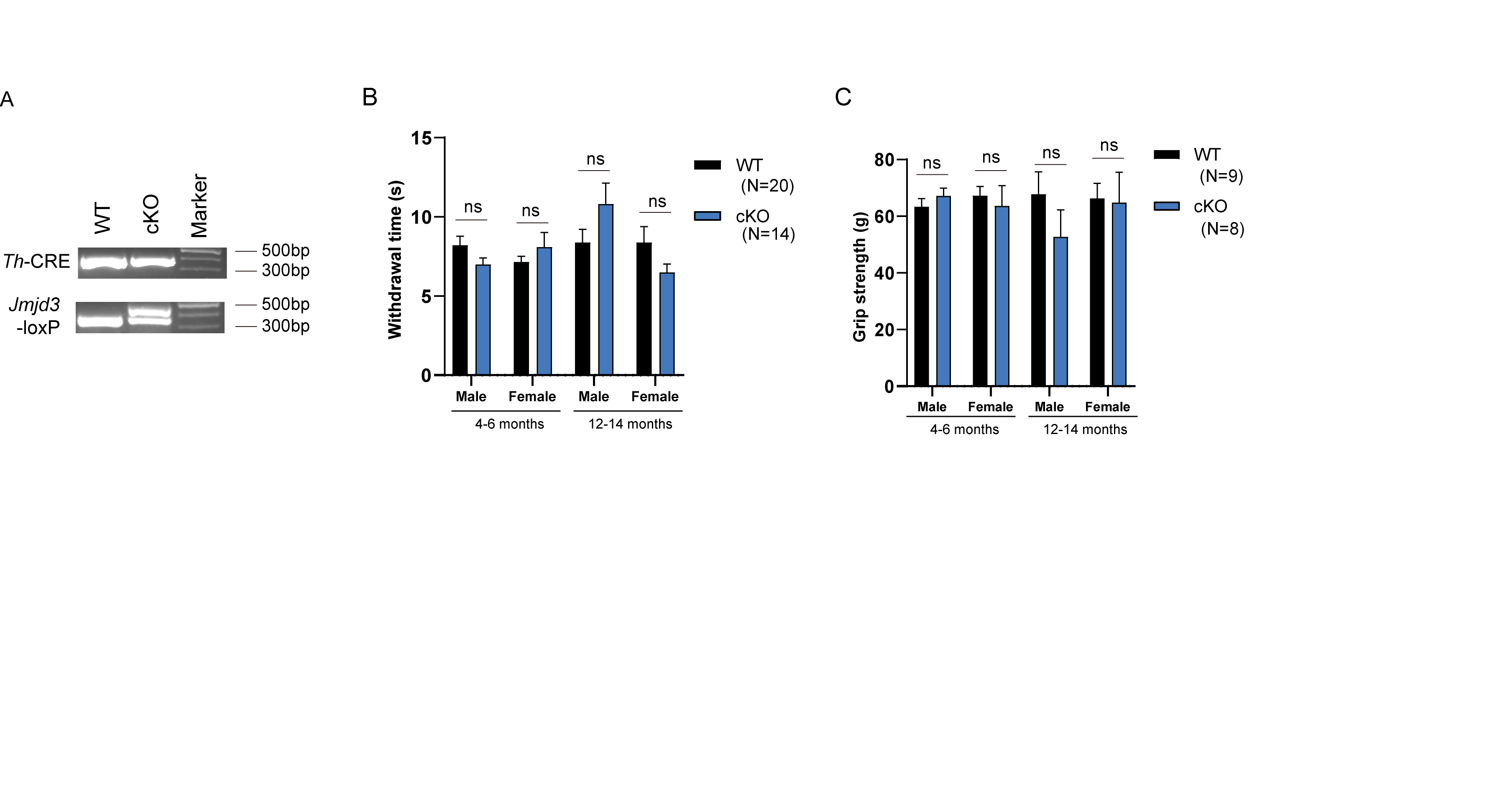

### Supplemental Figure 2

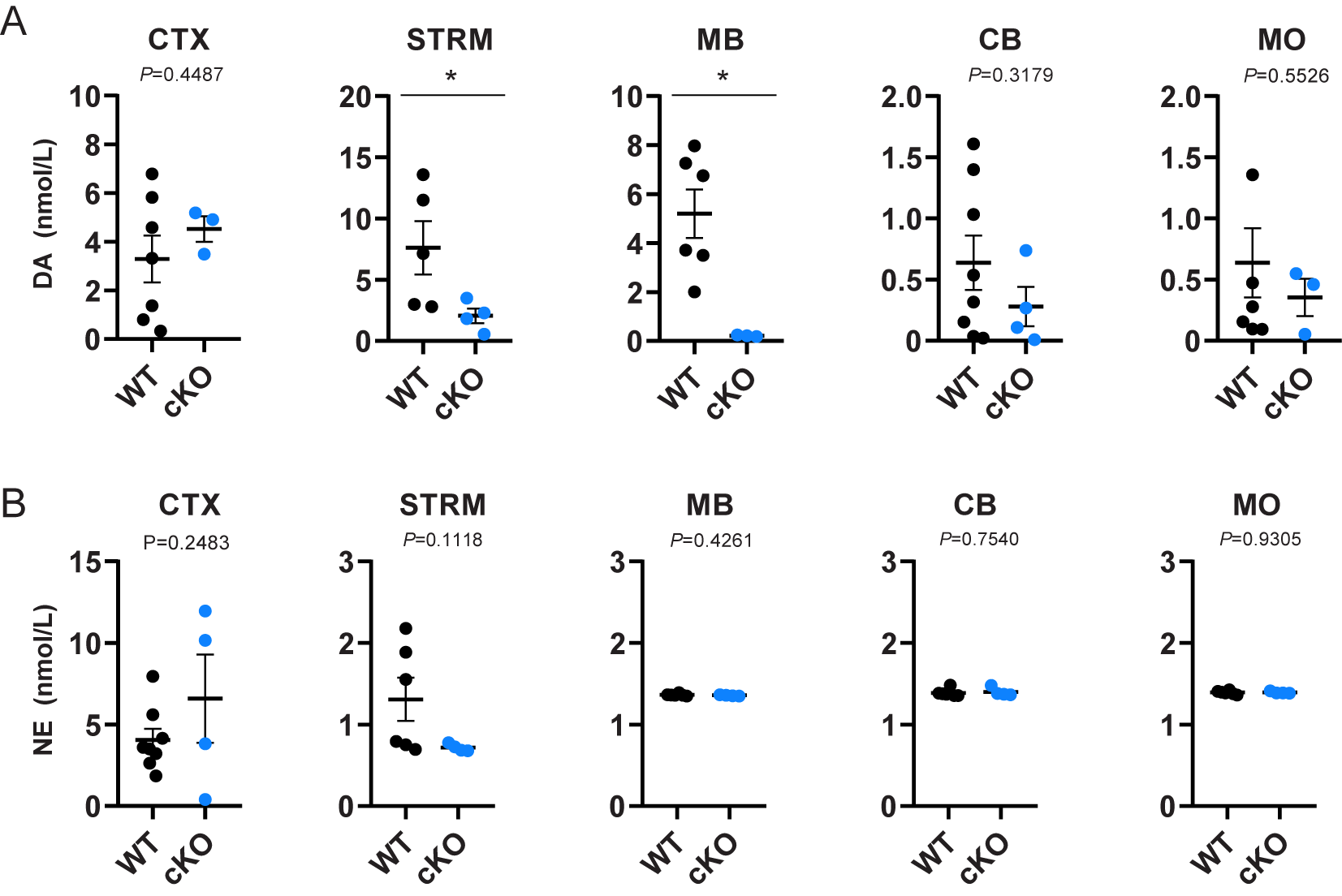

### Supplemental Figure 3

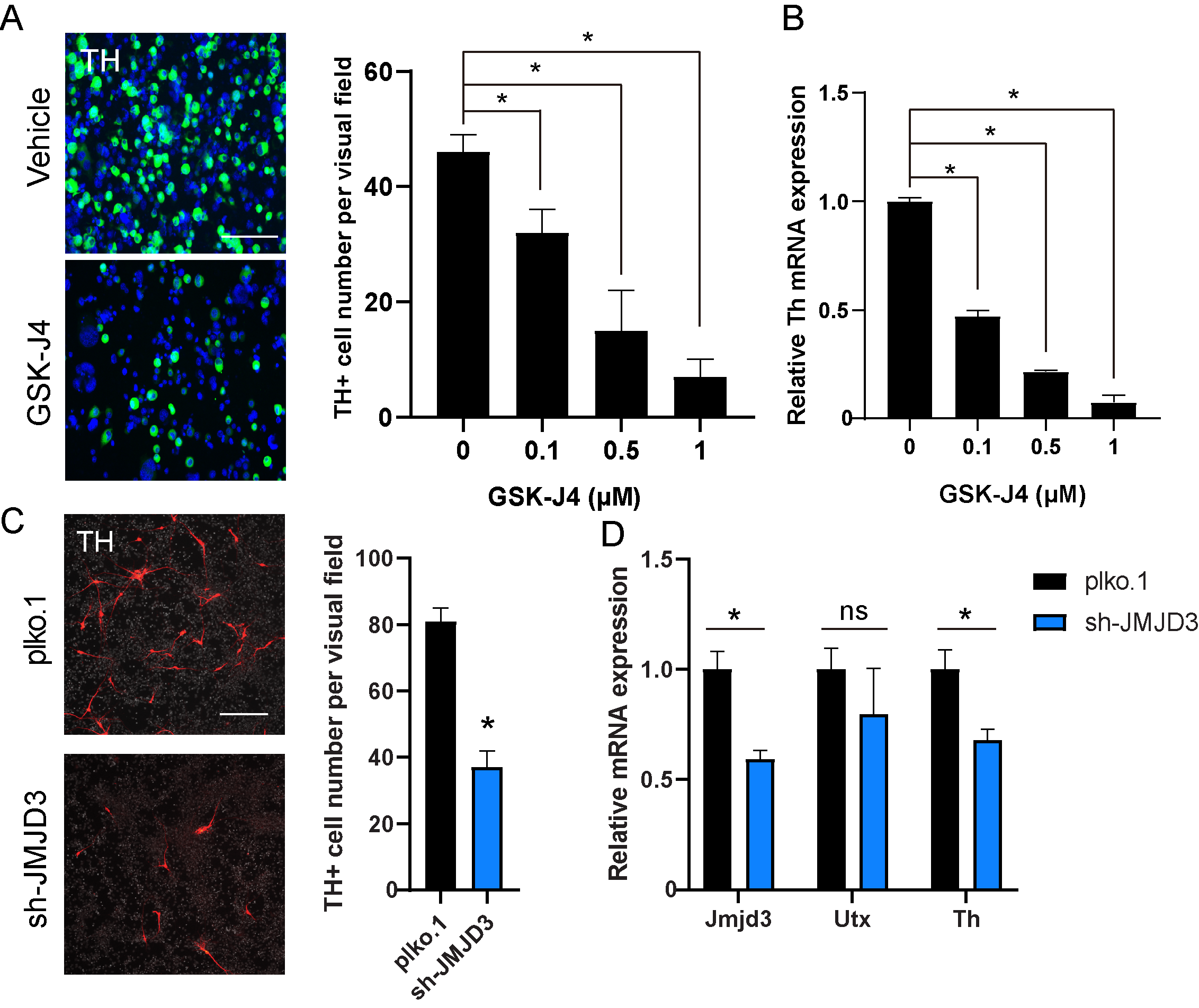
