## Supplemental Table 1 for "JMJD3 Deficiency in Midbrain Dopamine Neurons Disturbs Dopamine Synthesis and Aggravates Chronic Inflammatory Pain"

**Supplementary Table S1**

A. Primers for genotyping and quantitative PCR analysis

| Primer name | Forward | Reverse |
| --- | --- | --- |
| TH | GTCACGTCCCCAAGGTTCAT | GAGGAGGGTTTTGTACCCC |
| JMJD3 | CCCAGGCCCTGTGAGTAAAG | AGAGGCCAACGATTTGAGCA |
| UTX | CCTCCATTACCATCCGCCTC | AGAGTCCTGGCATAGGAGCA |
| GAPDH | TGTGAACGGATTTGGCCGTA | GTCTCGCTCCTGGAAGATGG |

| Primer name | Forward | Reverse |
| --- | --- | --- |
| CRE | TCGATGCAACGAGTGATGAG | TCCATGAGTGAACGAACCTG |
| JMJD3 loxP | GAGACCCCTGTTTGGTTACCAG | GAGGCAGAACCTGAGCAAGACC |

| Primer name | Forward | Reverse |
| --- | --- | --- |
| th1 | GCGACAGTGGATGCAATTAG | CCTCTTAAAGGCCAGGCTGA |
| th2 | GGGGACTTGAAGACATCCAA | CCCTAGACACGACATGAAGACA |
| th3 | TAGGGAGATGCCAAAGGCTA | AGTGTATGTGCTGGCACTGG |
| th4 | GCTCAGCATAAGTCCCCTGT | GTAAGGGCGCACTCAGTGAT |

| Primer name | Forward | Reverse |
| --- | --- | --- |
| Nurr1 | GCGGTGGGTCATTGTTTC | GCGCTCCGGTTCATTGTC |
| Nurr2 | GGGCACAGTGGCTTAAAAGT | CTCCTCTGCAAGTTCCAACC |
| Nurr3 | TGAATAAGACACGCGTCAGG | AGCCCCACTGTCCTTTCTTT |
| Nurr4 | CAGTGTCTTAGGGGCCAGAG | GAAGATCAGCTACTCTGCTGGA |

B. The parameters for multiple reaction monitoring (MRM)

| Compound | Precursor Ion | Product Ion | Fragmentor (V) | Collision Energy (eV) | Polarity | Quan or Qual |
| --- | --- | --- | --- | --- | --- | --- |
| Isoproterenol | 212.13 | 195.1 | 76 | 2 | Pos | Qual |
| Isoproterenol | 212.13 | 107 | 76 | 30 | Pos | Quan |
| HVA | 181.05 | 137.1 | 66 | 6 | Neg | Quan |
| 5-HT | 177.1 | 160 | 65 | 6 | Pos | Quan |
| 5-HT | 177.1 | 115 | 65 | 34 | Pos | Qual |
| NE | 174.1 | 156.1 | 118 | 10 | Neg | Quan |
| NE | 174.1 | 128.1 | 118 | 18 | Neg | Qual |
| DA | 154.09 | 137 | 70 | 6 | Pos | Quan |
| DA | 154.09 | 91 | 70 | 26 | Pos | Qual |
